# In *vivo* generation of BK and JC polyomavirus defective viral genomes in human urine samples associated with higher viral loads

**DOI:** 10.1101/2021.02.12.431053

**Authors:** Amin Addetia, Quynh Phung, Benjamin T. Bradley, Michelle Lin, Haiying Zhu, Hong Xie, Meei-Li Huang, Alexander L. Greninger

**Affiliations:** Department of Laboratory Medicine and Pathology, University of Washington, Seattle, Washington, USA; Molecular and Cellular Biology Graduate Program, University of Washington, Seattle, Washington, USA; Vaccine and Infectious Disease Division, Fred Hutchinson Cancer Research Center, Seattle, Washington, USA

## Abstract

Defective viral genomes (DVGs) are parasitic viral sequences containing point mutations, deletions, or duplications that might interfere with replication. DVGs are often associated with viral passage at high multiplicities of infection in culture systems but have been increasingly reported in clinical specimens. To date however, only RNA viruses have been shown to contain DVGs in clinical specimens. Here, using direct deep sequencing with multiple library preparation strategies and confirmatory ddPCR of urine samples taken from immunosuppressed individuals, we show clinical BKPyV and JCPyV strains contain widespread genomic rearrangements across multiple loci that likely interfere with viral replication. BKPyV DVGs were universally derived from type I subtype BKPyV. The presence of DVGs was associated with specimens containing higher viral loads but never reached clonality, consistent with a model of parasitized replication. These DVGs persisted during clinical infection as evidenced in two separate pairs of samples containing BK virus collected from the same individual up to 302 days apart. In a separate individual, we observed the generation of DVGs after a 57.5-fold increase in viral load. In summary, by extending the presence of DVGs in clinical specimens to DNA viruses, we demonstrate the ubiquity of DVGs in clinical virology.

**IMPORTANCE:** Defective viral genomes (DVGs) can have a significant impact on the production of infectious virus particles. DVGs have only been identified in cultured viruses passaged at high multiplicities of infection and RNA viruses collected from clinical specimens -- no DNA virus in the wild has been shown to contain DVGs. Here, we identified BK and JC polyomavirus DVGs in clinical urine specimens and demonstrated that these DVGs are more frequently identified in samples with higher viral loads. The strains containing DVGs had rearrangements throughout their genomes with the majority affecting genes required for viral replication. Longitudinal analysis showed these DVGs can persist during an infection, but do not reach clonality within the chronically infected host. Our identification of polyomavirus DVGs suggests these parasitic sequences exist across the many classes of viruses capable of causing human disease.

## INTRODUCTION

Defective viral genomes (DVGs) constitute a peculiar group of viral mutants incapable of replication. These DVGs contain point mutations, deletions, or duplications in their genomes (1). Many DVGs produce defective interfering particles (DIPs) that parasitize and interfere with viral replication. DVGs/DIPs are thought to contribute to chronic viral infections by promoting survival of the infected host cells or reducing the infectious viral load by suppressing the number of replication-competent virions produced during active infection (2). Increasingly, DVGs are being investigated for their potential as antiviral strategies (1, 3).

Two major types of DVGs have been described for RNA viruses (1). Deletion-type DVGs are truncated versions of the non-defective viral genome that occur when the polymerase skips part of the genome during replication. Deletion DVGs tend to possess identical terminal sequences while missing several or all essential genes required for self-propagation (4, 5). The second type of DVGs is the copy-back DVGs. Copy-back and the related snap-back DVGs are rearranged genomes consisting of an authentic terminus followed immediately with an inverted repeat of some or all of that sequence (6). Copy-back DVGs have been reported in many negative-sense RNA viruses, mainly in the *Paramyxoviridae* family, and are predicted to be the products from the reattachment of RNA-dependent RNA polymerase (RdRp) to the nascent strand, thus copying back the end of the genome (7).

DVGs have frequently been identified in cell culture systems, especially when viral stocks are passaged at high multiplicities of infections, and are increasingly being identified in clinical specimens (2, 8–12). These DVGs contain deletions of variable lengths and across various regions in the genome, but generally retain the origin of replication (13). Cultured BK and JC polyomavirus have both been shown to contain deletions spanning the large T antigen (14, 15). To date, only single-stranded RNA viruses have been demonstrated to form DVGs in clinical specimens (2, 16–18). However, the significance and role of DVG during clinical infection are unclear.

Polyomaviruses are non-enveloped, double-stranded DNA viruses with a small, circular genome of approximately 5000 base pairs (19). Among the 102 identified species, BK polyomavirus (BKPyV) and JC polyomavirus (JCPyV) are the two most commonly known to infect humans (20). BKPyV and JCPyV are highly prevalent in the population with most individuals initially getting infected in early childhood and maintaining a lifelong infection thereafter (21, 22). The infections of BKPyV, also known as Human polyomavirus 1, are generally asymptomatic in immunocompetent individuals, but can cause nephropathy and allograft failure in individuals receiving a renal transplant (23, 24). In individuals with hemorrhagic cystitis, BKPyV can be found at very high titers in urine (> 10^8^ copies/mL of urine) (8). JCPyV, also known as Human polyomavirus 2, is most well-known for its association with progressive multifocal leukoencephalopathy (PML) in immunosuppressed individuals, and can cause many novel neurological disorders such as JC virus granule cell neuropathy and JC virus encephalopathy (9). JCPyV can be found in cerebrospinal fluid and urine of individuals with PML at values ranging from 10^2^ to 10^8^ copies/mL (10).

While performing genome recovery of polyomaviruses for urine specimens sent to our clinical laboratory, we noticed many large-scale genomic rearrangements in BKPyV and JCPyV. These rearrangements were recovered independently of the sequencing strategy used and were found across different viral lineages. Polyomavirus DVGs were found more frequently in those specimens that were specifically associated with high viral load specimens but never reached clonality, consistent with a model of defective interfering replication.

## RESULTS

### Defective polyomaviruses genomes in clinical urine samples

We performed metagenomic shotgun sequencing on DNA extracted from 44 BKPyV-positive clinical urine samples collected from 38 individuals with a median viral load of 1.30 × 10^8^ copies/mL (range: 6.3 × 10^4^ - 4.6 × 10^10^ copies/mL) and 2 JCPyV-positive clinical urine specimens (Table S1). Of the 46 samples, we recovered 39 complete BKPyV genomes and 11 complete JCPyV genomes and found BKPyV and JCPyV co-infections in 11 of these samples (Figure S1). We identified co-occurrence of common uropathogenic bacteria present at a frequency greater than 10 reads per million (RPM) in 31 of the 46 polyomavirus-positive samples (Table S2). Reads corresponding to *Candida* spp. at a frequency greater than 10 RPM were identified in 9 of the 46 samples.

In 13 of the BKPyV-positive samples, we identified large rearrangements or deletions constituting DVGs in BKPyV or JCPyV that were supported by 10 or more sequencing reads and included samples with a sum total frequency of 10% or greater of these rearrangements (Figure 1, Table S3). Twelve of the 13 samples with rearrangements were identified in BKPyV genomes collected from 11 different individuals, while the remaining sample was identified in a JCPyV genome.

**Figure 1.**
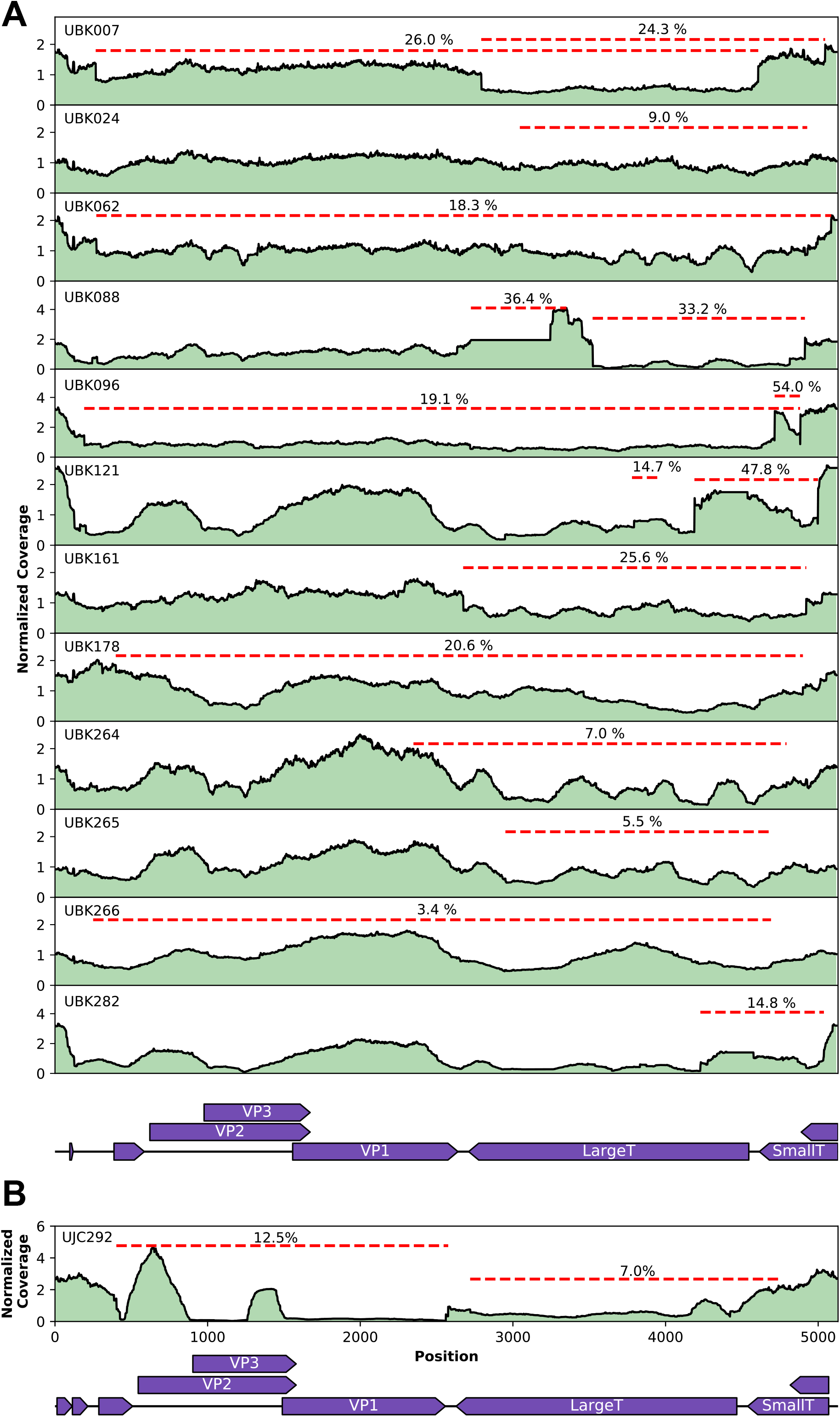
Coverage maps of defective BKPyV genomes (**A**) and JCPyV genome (**B**) observed in shotgun sequencing data of 13 polyomavirus positive specimens. Normalized coverage is plotted after mapping to each sample’s consensus viral genome sequence. Representative junction reads are depicted in red dashes along with the percentage of the junction reads relative to the maximum coverage.

The nucleotides present in the junctions in the BKPyV strains were separated by a range of 4 to 2491 nucleotides (median: 628 nt). Notably, multiple unique junctions (median: 15; range: 2-40) were identified in each DVG-containing sample. All 12 BKPyV strains had junctions including the large T antigen and in 10 of the strains, the most abundant deletion or rearrangement involved the large T antigen (Table S3A). The most abundant junction in 11 of the BKPyV strains were internal deletions and, correspondingly, these DVGs were classified as deletion-type. In the final strain, UBK096, the most abundant junction was an inversion rearrangement. In the 1 JCPyV strain with detectable junctions, 15 unique junctions, ranging from 285-2302 nucleotides in length (median: 1917 nt), were identified (Table S3B). The most abundant junction was a rearrangement that spanned all three capsid proteins, VP1, VP2, and VP3. Other DVGs in the JCPyV present in specimen UBK292 were classified as deletion-type.

We confirmed the observed junctions were not an artefact of our library preparation protocol by performing a second method for generating sequencing libraries on a subset of DVG-containing and DVG-negative samples. Identical junctions were found in the DVG-containing samples and no new DVGs were found in DVG-negative samples using the different library preparation (Figure S2). We further confirmed select junctions using specific PCR and Sanger sequencing (NCBI BioProject PRJNA657423).

### An elevated VP1-to-large T antigen ratio is observed in BKPyV strains containing defective viral genomes and having high viral loads

We have previously shown that digital droplet PCR (ddPCR) is a cost-effective method to detect and confirm copy number alterations in cultured BKPyV and JCPyV (15). Since many DVGs are also associated with copy number alterations, to confirm our sequence-based results from clinical specimens, we performed ddPCR targeting the VP1 and large T antigen on the 12 BKPyV strains containing DVGs by deep sequencing and 22 BKPyV strains without DVGs by deep sequencing.

Quantitative analysis with ddPCR assay showed that the VP1-to-large T antigen copy number ratio was significantly higher in those strains with DVGs (mean: 1.69; range: 0.99 – 3.57) than those with intact genomes (mean: 1.03; range: 0.91 – 1.31) (p=0.0002, Kruskal-Wallis test, Figure 2A). Of note, two strains with DVGs (UBK265/UBK266) had a comparably low level of rearrangements (3-5%) by sequencing at the exact loci interrogated by the ddPCR and had a ddPCR VP1/large T antigen ratio equivalent to 1. A VP1/large T antigen ratio cutoff of 1.25 separated the remaining 10 DVG-containing strains confirmed by sequencing from all strains without DVGs except UBK292.

**Figure 2.**
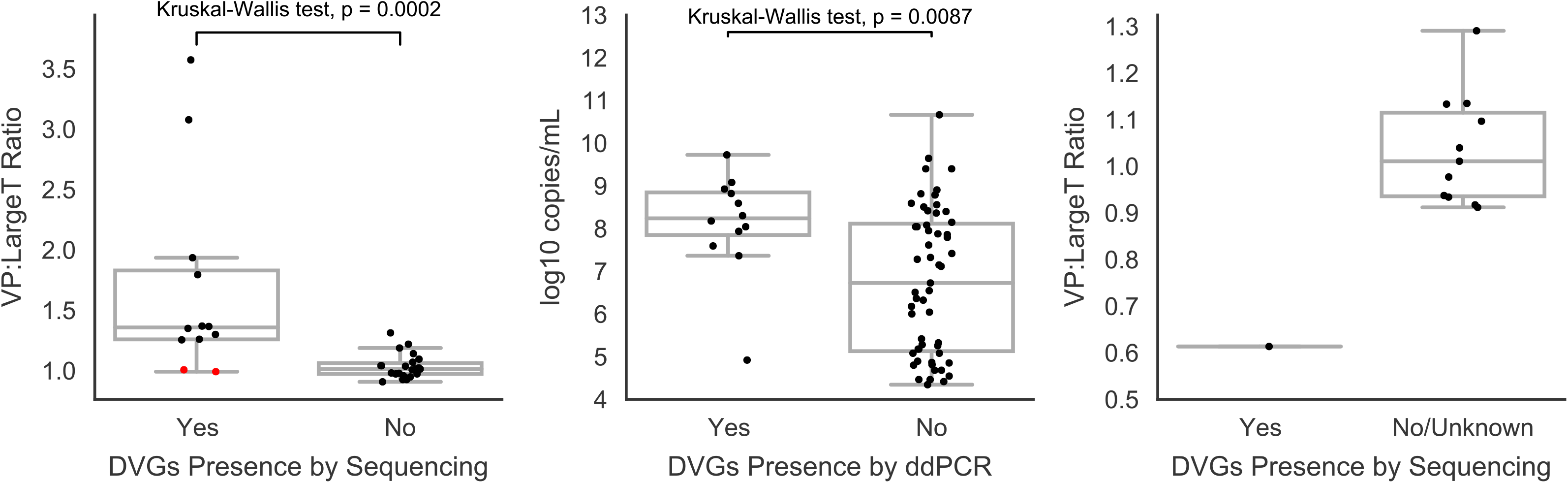
Confirmation of defective viral genomes using droplet digital PCR. The copy number ratio for VP1 and Large T antigen is plotted for (**A**) 34 BKPyV isolates with and without DVGs determined by sequencing. Quartiles for each group are plotted in a box-and-whiskers plot and error bars are 1.5-fold the interquartile range. Red dots indicate two strains (UBK265/UBK266) with DVGs that had a comparably low level of rearrangements (3-5%) at the target loci as determined by sequencing, consistent with their equivalent copy number measured by ddPCR. Statistical comparison was performed via Kruskal-Wallis test. (**B**) Log_10_ copies/mL viral load values in 67 BKPyV-positive specimens are displayed. The presence of DVGs was determined by ddPCR using a VP1/largeT antigen ratio cutoff of 1.25. Statistical comparison was performed via Kruskal-Wallis test. (**C**) ddPCR VP1/Large T antigen copy number ratios for 12 JCPyV-positive specimens are depicted. One JCPyV specimen contained a sequence confirmed DVG while 7 of the 11 other specimens lacked DVGs by sequencing and the remaining had unknown DVG status.

We next used this ddPCR and established ratio cutoff to screen 33 additional BKPyV positive specimens for which high coverage genomes could not be recovered by shotgun sequencing due to low viral load. Notably, just one of the 33 low viral load samples had a VP1/Large T antigen ratio greater than 1.25, consistent with low viral load infrequently containing DVGs. Furthermore, this analysis showed that specimens that tested positive for copy number alterations by ddPCR (VP1/Large T antigen ratio > 1.25) had a greater than 33-fold higher median viral load than those specimens that had a normal copy number ratio (median 1.75 × 10^8^ copies/mL versus 5.30 × 10^6^ copies/mL, p=0.0087, Kruskal-Wallis test) (Figure 2B).

We next performed ddPCR targeting the VP1 and large T antigen on the JCPyV strain containing DVGs and 11 JCPyV-positive urine specimens, 7 of which did not contain DVGs by deep sequencing. The 11 JCPyV-positive strains had a mean VP1/Large T antigen copy number ratio of 1.03 (range: 0.91 – 1.29), while the strain with the DVGs had a ratio of 0.61 (Figure 2C), consistent with a copy number ratio of 0.58 by sequencing at the targeted loci.

### Phylogenetic analysis of BKPyV and JCPyV genomes reveals strains containing defective viral genomes belong to multiple subgroups and are stable across time

We next assessed the genetic relatedness of the BKPyV strains that produced DVGs by performing a phylogenetic analysis with 106 representative BKPyV genomes and the 39 total BKPyV genomes recovered in this study (Figure 3A). Of the 39 strains, 37 belonged to subtype I, 1 belonged to subtype III, and 1 belonged to subtype IV. These observations are consistent with subtype I being the most prevalent subtype in the United States and subtypes II-IV being less frequently detected in North America (25). 18 of the 37 subtype I strains belonged to subgroup Ia, 8 belonged to subgroup Ib-1, and 11 belonged to subgroup Ib-2. The 12 strains containing DVGs all belonged to subtype I. Ten of these strains belonged to subgroup Ia, 1 belonged to subgroup Ib-1, and 1 belonged to subgroup Ib-2.

**Figure 3.**
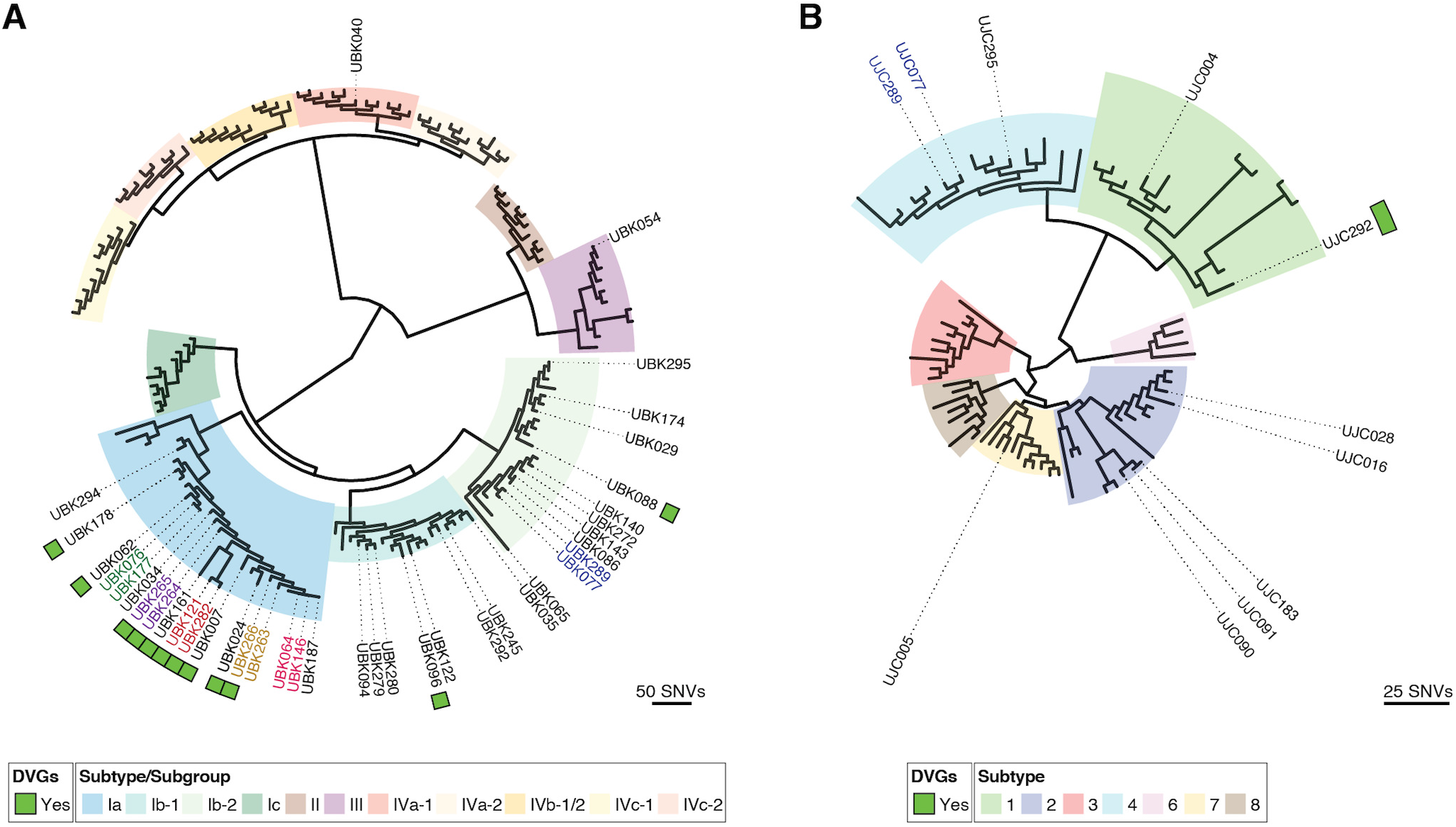
**(A)** Phylogenetic tree of 39 BKPyV consensus genomes recovered in this study along with 106 representative BKPyV genomes covering all 11 subtypes. BKPyV samples containing DVGs are indicated by a green square and those samples longitudinally collected from the same individual are highlighted through coloring of the sample labels. (**B**) Phylogenetic tree of the 11 JCPyV genomes recovered in this study with 106 JCPyV covering 7 JCPyV subtypes. The samples containing DVGs are indicated with a green square.

Of the 39 BKPyV genomes recovered in this study, 6 pairs of isolates were collected from the same individuals (UBK064/UBK146; UBK077/UBK289; UBK076/UBK177; UBK121/UBK282; UBK264/UBK265; UBK263 and UBK266), allowing us to look at longitudinal evolution of BKPyV DVGs. The first three of these pairs (UBK064/UBK146; UBK077/UBK289; UBK076/UBK177) did not contain any DVGs and were genetically identical across a longitudinal sampling time of 7, 487, and 21 days, respectively. Interestingly, in one pair collected 159 days apart, the first isolate UBK263 only contained intact genomes while the second isolate UBK266 contained DVGs. Only 1 consensus single nucleotides variant (SNV) was recovered between these samples, which resulted in a coding change (Glu43Lys) in the agnoprotein. Notably the BKPyV viral load increased by 57.5x between UBK263 and UBK266, suggesting the increase in viral load may have contributed to the formation of DVGs. In the final two pairs (UBK121/UBK282 and UBK264/UBK265), both strains contained DVGs. UBK121 and UBK282 were collected 302 days apart and differed by 2 SNVs as well as a 39bp insertion at the origin. One of the SNVs resulted in a coding change, Lys556Arg, in the large T antigen. UBK264 and UBK265 were collected 41 days apart and differed by 2 SNVs, both of which resulted in coding changes in the agnoprotein (Gln8Arg and Glu43Lys).

We then assessed the genetic relatedness of the 11 JCPyV strains sequenced in this study by performing a phylogenetic analysis with 64 representative JCPyV genomes (Figure 3B). Two of the strains belonged to subtype 1, 5 belonged to subtype 2, 3 belonged to subtype 4, and 1 belonged to subtype 7. The one JCPyV strain containing DVGs belonged to subtype 1.

### Persons with DVG-containing viruses more likely to have changes to immunosuppressive regimens

Clinical information was available for 32 individuals with BK viruria including 22 individuals without any samples containing DVGs and 10 individuals with DVGs identified in at least one clinical sample (Table 1). Both the DVG-negative and DVG-containing groups were similar in age (mean 55+/− 16.3 v. 53+/− 19.9 years) and proportion of females (54.5% and 50.0%) respectively. The clinical setting under which all individuals developed BK viruria was related to immunosuppression secondary to organ transplant. Renal transplant was the most common indication for both groups with peripheral blood stem cell and multiorgan (kidney and pancreas or kidney and liver) transplants representing a smaller percentage. Comparison of immunosuppressive regimens between DVG groups demonstrated that DVG-containing individuals more often had their dose of mycophenolate stopped or changed to leflunomide. Changes in therapy for the DVG-containing group are perhaps related to their higher viral loads.

**Table 1A/B.**
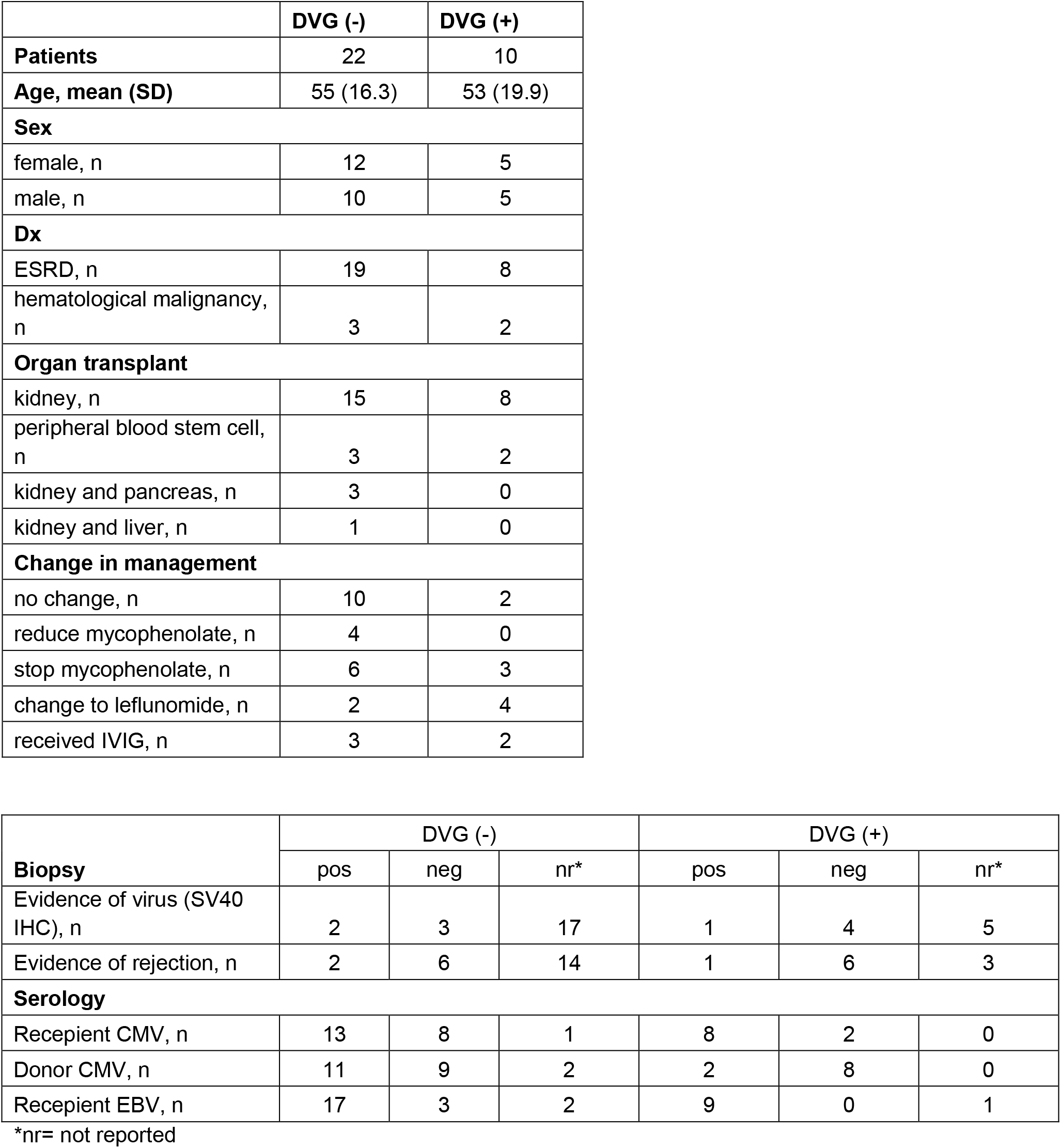
Clinical characteristics of the individuals in this study.

DVG-containing individuals were more likely to undergo renal biopsy (7 of 10 vs. 8 of 22). However, histologic evidence of kidney injury or immunohistochemical staining for the SV40 large T antigen was rarely seen (1 and 4 individuals, respectively). Pre-transplant serologies demonstrated the majority of individuals in both groups to be CMV and EBV positive. In the DVG-containing group, individuals more often received organs from donors who were CMV-negative.

## DISCUSSION

Here, we report the presence of polyomavirus defective genome segments in clinical specimens and observe the diversity of DVG populations naturally generated across individuals and within a host over the course of infection. Specifically, by deep sequencing, we identified multiple BKPyV strains and one JCPyV strain that contained large deletions or rearrangement junctions in their genomes. Although these junctions have been observed in all six gene segments, most were present in the large T antigen, which is consistent with reports from other viruses in which DVGs most commonly occur in replication-related genes (11). These observations also are consistent with the characteristic features of the polyomavirus DVGs found in culture: major internal deletions with retention of regions that are essential for genome packaging (2, 11, 18).

DVG formation is thought to be a form of viral parasitism, which occurs when multiple viruses infect the same cell (2, 3, 11, 18). Our finding that DVGs were more likely to be present when polyomavirus was present at high copy numbers is consistent with this model. A dramatic increase in viral load was also temporally associated with the only longitudinal specimens in which DVGs newly arose. In addition, the bidirectional replication mechanism of polyomaviruses combined with decatenation of linked circular genomes would provide several opportunities for recombination and generation of deletions and other large-scale rearrangements (12, 26, 27).

We also examined the evolution of BKPyV in a small subset of individuals that had longitudinal specimens available. The recovery of 1-2 SNVs in specimens taken from persons with longitudinal samples spaced between 1 month −1 year apart is consistent with the relatively high intra-host evolutionary rate of polyomaviruses, previously measured at 4.9 × 10^−4^– 1.2 × 10^−3^ substitutions per site per year (28). Given that we only detected one newly generated DVG in this study, more work is required to uncover the determinants and rate of DVG generation in polyomaviruses.

One main limitation of our sequencing approach was the use of short-read sequencing, which may have affected our measurements of DVGs in each specimen and did not allow us to link rearrangements across the multiple populations of virus present within the viral population. Recent work in polyomaviruses has demonstrated the promise of long-read sequencing to resolve complex populations of polyomaviruses (29, 30). However, the limited total DNA contained in these samples complicates the use of long-read approaches given the large-scale rearrangements recovered and requirement for multiple cycles of PCR. We also quantitated rearrangements as a percent of maximum coverage which may bias quantitation depending on local variations in copy number. We bulwarked our sequencing approach by using multiple library preparations of the same sample to illustrate that deletions and rearrangements recovered were not due to library generation.

Of note, it has yet to be established whether these DVGs identified from the clinical specimens interfere with the replication of the full-genome virus *in vitro* and the role of DVGs in natural polyomavirus infections has yet to be addressed. The deletions and rearrangements recovered in our study were generally quite large and affected proteins and domains required for viral replication, consistent with their defective nomenclature. Further studies focusing on the function of specific rearrangements found in polyomavirus DVGs through reverse genetics and cell culture-based approaches are required to help understand their role in replication interference.

In conclusion, by demonstrating the existence of polyomavirus DVGs in human specimens, we extend this broad property of viral evolution to DNA viruses as they exist in human hosts. The polyomaviruses containing DVGs recovered in this study generally had significantly higher copy number of capsid genes to replication genes. Further work is required to determine whether these genetic copy alterations result in differences in protein levels in human samples that could potentially affect pharmacodynamic properties and efficacy profiles of capsid directed therapies for polyomaviruses.

## METHODS

### Clinical Testing and Next-generation Sequencing

This work was approved by the University of Washington Institutional Review Board (STUDY00000408). Urine samples were collected from individuals suspected to have a BKPyV or JCPyV infection and clinical testing was performed at the University of Washington Virology Laboratory using the previously described quantitative PCR assays (14, 31).

BKPyV-or JCPyV-positive urine samples were first filtered using a 0.22 μM filter and DNA was extracted from these samples using the Quick-DNA Viral Kit (Zymo). Sequencing libraries were constructed from 2 μL of DNA using the Nextera XT Kit (Illumina) and cleaned with 0.6x volumes of Ampure XP beads (Beckman Coulter). For four samples, a second sequencing library was created using the KAPA HyperPrep Plus kit (Roche) and purified with 0.8x volumes of AMPure XP beads. The resulting libraries were then sequenced on 1×192 bp or 2×300 bp Illumina MiSeq runs.

### Identification of Defective Viral Genomes

Sequencing reads were adapter-and quality-trimmed using Trimmomatic v0.38 (32) and mapped to the BKPyV (NC_001538.1) or JCPyV (NC_001699.1) reference genomes. Consensus genomes were manually called and sequencing reads were then mapped to the respective consensus genome in Geneious Prime (33) with the structural variant, insertion, and deletion detection setting enabled. Because of the high proportion of small deletions contained around the origin and upstream of the agnoprotein, those junctions entirely contained between nucleotides 0-300 were discarded and not included in downstream further analysis. Each of the junctions reported here were supported by a minimum of 10 sequencing reads. In addition, we confirmed the presence of the highest abundance junction identified in each sample using DI-tector (34).

Next, we confirmed the presence of these junctions in a subset of these samples through PCR and Sanger sequencing. PCR amplification was performed using the CloneAmp HiFi Premix (Takara) and the primers listed in Table S4 under the following conditions: hold at 98°C for 2 minutes, followed by 35 cycles of 98°C for 10 seconds, 61°C for 15 seconds and 72°C for 30 seconds, followed by a final extension at 72°C for 5 minutes. PCR products were run on a 1.5% agarose gel and extracted using the QIAquick Gel Extraction Kit (Qiagen). Sanger sequencing was performed with the primers listed in Table S1 by Genewiz, Inc.

### Droplet digital PCR Detection of Defective Viral Genomes

ddPCR was performed using the Bio-Rad QX100 system (Bio-Rad, Hercules, CA, USA) and QuantaSoft for data analysis. There are two sets of primers and probes targeting the VP1 and Large T antigen regions, respectively (Table S4). Each reaction was performed with BioRad ddPCR Supermix for Probes with final concentration of primers at 900nM and probes at 250nM and 25 units of HindIII (New England Biolabs). Plasmid BK Dunlop and JC Mad-1 were gifts from Peter Howley (Addgene plasmids # 25466 and #25626) and were used as positive controls for 1:1 VP1/Large T-antigen copy number. After droplet generation, droplets were transferred to a 96-well PCR plate and amplified on a 2720 Thermal Cycler (Applied Biosystems) with the following thermocycling parameters: 94°C for 10 min, followed by 40 cycles of 94°C for 30 s and 60°C for 1 min, and 98°C hold for 10 min. After the thermal cycling, the plate was transferred to a droplet reader. The QuantaSoft software was used for data analysis.

### Phylogenetic and metagenomic analysis

We downloaded all 510 complete BKPyV genomes from NCBI GenBank (accessed August 16, 2020) and removed any duplicate genomes from this dataset to obtain 402 unique BKPyV genomes. These 402 BKPyV genomes were then classified into the 11 VP1 sequence subtypes and subgroups with BKTyper 1.0 (35). We next randomly selected 10 genomes from each of these subtypes or subgroups for inclusion in our phylogenetic analysis. The 106 representative BKPyV genomes and the 39 complete BKPyV genomes obtained in this study were aligned with MAFFT v7.429 (36). A phylogenetic tree was generated from this alignment using RaxML v8.2.11 (37) and visualized with ggtree (38).

To perform the JCPyV phylogenetic tree, we downloaded all 696 complete JCPyV genomes from NCBI GenBank (accessed January 16, 2021) and removed any duplicate genomes to obtain 565 unique JCPyV genomes. These genomes were then classified into the previously defined JCPyV subtypes (39) based on their VP1 sequence using a custom Python script. We then randomly selected genomes from each subtype. An alignment and phylogenetic tree were generated as described above with these 64 representative JCPyV genomes and the 11 JCPyV recovered in this study.

Metagenomic analysis was performed as previously described (40), using the metagenomic classifier CLOMP (https://github.com/rcs333/CLOMP). Counts were normalized between samples and classifications were expressed as reads per million (RPM). Taxonomic classifications of each read were assigned to the most specific NCBI taxonomy ID possible and visualized with ggplot. Any reads assigned to environmental or artificial sequences were discarded. Reads that matched equally well to two or more different domains were categorized as “Unclassified.”

## Data Availability

Sequencing reads and consensus genomes are available under NCBI BioProject PRJNA657423 and accession numbers are additionally listed in Table S5.

